# Sex and pubertal status influence dendritic spine density onto frontal corticostriatal projection neurons

**DOI:** 10.1101/787408

**Authors:** Kristen Delevich, Nana J. Okada, Ameet Rahane, Zicheng Zhang, Christopher D. Hall, Linda Wilbrecht

## Abstract

In humans, nonhuman primates, and rodents, the frontal cortices exhibit grey matter thinning and dendritic spine pruning that extends late into adolescence. This protracted maturation is believed to support higher cognition but may also confer psychiatric vulnerability during adolescence. Currently, little is known about how different cell types in the frontal cortex mature or whether puberty plays a role. Here, we used mice to characterize the spatial topography and adolescent development of cross-corticostriatal (cSTR) neurons that project to the dorsomedial striatum (DMS). We found that apical spine density on cSTR neurons in the medial prefrontal cortex decreased significantly between late juvenile (P29) and young adult time points (P60), with females exhibiting higher spine density than males at both ages. Adult males castrated prior to puberty onset had higher spine density compared to sham controls. Adult females ovariectomized before puberty onset showed greater variance in spine density measures on cSTR cells compared to controls, but their mean spine density did not significantly differ from sham controls. Our findings reveal that these cSTR neurons, a subtype of the broader class of intratelencephalic-type neurons, exhibit significant sex differences and suggest that spine pruning on cSTR neurons is regulated by puberty in males.

---

Adolescence is the developmental transition between juvenile and adulthood that is accompanied by significant changes in brain structure and function (Spear, 2000). While puberty typically marks the onset of the adolescent period, it is still unclear whether adolescence-associated brain changes are driven by pubertal development. Across species, adolescence is characterized by newfound independence and changing social roles, and while the richness of human social and cultural experience cannot be captured in rodent models, the functional consequences of puberty can be more easily translated across species.

Structural imaging studies in humans have shown that frontal cortical gray matter undergoes significant thinning during adolescence (Sowell et al., 1999), and postmortem studies provide evidence that the reduction in gray matter volume may be attributed, in part, to synapse elimination that occurs during this time (Huttenlocher, 1979; Rakic et al., 1986; Bourgeois et al., 1994; Anderson et al., 1995; Glantz et al., 2007; Petanjek et al., 2011). Changes in frontal gray matter overlaps with pubertal milestones (Giedd et al., 1999), raising the question of whether the pubertal increase in steroid sex hormones drives synaptic pruning during adolescence. Recently, human neuroimaging studies have begun to parse the effects of age and pubertal status on brain development (Neufang et al., 2009; Peper et al., 2009; Paus et al., 2010; Bramen et al., 2011). Studies have found that testosterone is associated with decreases in cortical thickness in postpubertal males (Nguyen et al., 2013) and gray matter volume in the frontal lobes (Koolschijn et al. 2014). Another found that pubertal tempo (the rate of change in Tanner staging) predicted the rate of change in cortical thickness in some regions (Herting et al. 2015). Dendritic spines in the frontal cortex have also been shown to prune across adolescence in rats (Koss et al., 2014) and mice (Holtmaat et al., 2005; Gourley et al., 2012; Johnson et al., 2016; Boivin et al., 2018). Similarly, rats show a decrease in overall synaptic density in the medial PFC (mPFC) across adolescence, and in both males and females spine density was lower in rats with more advanced pubertal status when compared with same age pre-pubertal controls (Drzewiecki et al., 2016).

While these data strongly suggest that puberty influences frontal cortex synapse elimination, less is known about the specific cell types and circuits involved. In the cortex, two major excitatory pyramidal neuron classes include intratelencephalic IT-type neurons, which mediate cortico-cortical communication, and pyramidal tract PT-type neurons that project outside the telencephalon to the brainstem and spinal cord (Shepherd, 2013; Harris and Shepherd, 2015). IT-type and PT-type neurons are intermingled within layer (L)5, but exhibit distinct morphological and electrophysiological properties and target different downstream brain areas (Cowan and Wilson, 1994; Reiner et al., 2003; Gerfen et al., 2013; Naka and Adesnik, 2016; Baker et al., 2018). Notably, while both IT-type and PT-type neurons project to the striatum, only IT-type neurons cross the corpus callosum to innervate contralateral striatum, which we refer to as cross-corticostriatal (cSTR) neurons (Shepherd, 2013; Harris and Shepherd, 2015). Recent work in rodent models suggests that prefrontal IT-type neurons may exhibit a more protracted maturation than PT-type neurons. Data from mouse mPFC suggests that synaptic inhibition onto L2/3 and 5 IT-type neurons is dynamic during the adolescent period between postnatal day (P)25 and 45, whereas inhibition onto L5 PT-type neurons is stable (Vandenberg et al., 2015; Piekarski et al., 2017a). Furthermore, ovarian hormones at puberty appear to play a causal role in the maturation of inhibitory synaptic transmission onto IT-type neurons in female mice (Piekarski et al. 2017a). In L5 PT-type neurons labeled by the YFP-H transgenic mouse line (Porrero et al., 2010), data suggests that ovarian hormones at puberty do not influence spine pruning in female mice (Boivin et al., 2018). We have therefore developed a hypothesis that IT-type neurons may play an important role in adolescent reorganization of the brain, particularly aspects that are regulated by puberty onset (Delevich et al., 2019b).

Given that many aspects of goal-directed behavior mature across adolescence (Huizinga et al., 2006; Johnson and Wilbrecht, 2011; Blakemore and Robbins, 2012), and that cSTR projections to the DMS have been implicated in goal-directed learning (Hart et al., 2018a; Hart et al., 2018b), we were motivated to compare apical spine density onto DMS-projecting cSTR neurons between juvenile and adult time points. Significant remodeling of excitatory inputs onto this population of cSTR neurons across the adolescent transition could potentially support the adolescent maturation of goal-directed behavior (Naneix et al., 2012; DePasque and Galvan, 2017). Our goals for the current study were to: 1) characterize frontal cSTR neurons projections and their intrinsic properties across adolescence, 2) determine if prefrontal cSTR IT-type neurons exhibit spine pruning during adolescence in male and female mice and 3) determine whether adolescent spine pruning onto cSTR IT-type neurons is sensitive to prepubertal gonadectomy.

## MATERIALS & METHODS

### Animals

Male and female C57BL/6NCR mice (Charles River) were bred in-house. All mice were weaned on postnatal day (P)21 and housed in groups of 2–3 same-sex siblings on a 12:12 hr reversed light:dark cycle (lights on at 2200h). All animals were given access to food and water ad libitum. For the developmental comparison, our juvenile time point was postnatal day (P)29 (7 females, 6 males) and our adult age was P60 (6 females, 4 males), an age range spanning the adolescent period (Tirelli et al., 2003). For our puberty comparison, P60 was used as the age for both the sham (5 females, 4 males) and GDX (6 females, 7 males) groups. All procedures were approved by the Animal Care and Use Committee of the University of California, Berkeley and conformed to principles outlined by the NIH Guide for the Care and Use of Laboratory Animals.

### Stereotaxic injections

Male and female mice (P21 or P52) were deeply anesthetized with 5% isoflurane (vol/vol) in oxygen and placed into a stereotactic frame (Kopf Instruments; Tujunga, CA) upon a heating pad. Anesthesia was maintained at 1-2% isoflurane during surgery. An incision was made along the midline of the scalp and small burr holes were drilled over each injection site. Virus or tracer was delivered via microinjection using a Nanoject II injector (Drummond Scientific Company; Broomall, PA). For spine imaging experiments, 50 nL pAAV-CAG-GFP (Addgene item #37825-AAVrg) diluted 1:5 in saline (starting titer ≥ 7×10^12^ vg/mL) was delivered unilaterally to the dorsomedial striatum (DMS) (coordinates relative to Bregma: A/P: +.9mm, M/L: +1.4mm, D/V: +3.0 mm for adults, +2.7 mm for juveniles). For electrophysiology, dual labeling (Figure 1), and input mapping (Figure 2) experiments, 100-200 nL of 1:5 dilute retrograde AAV virus [pAAV-CAG-tdTomato or pAAV-CAG-tdTomato] was delivered unilaterally to the DMS and/or pons (coordinates relative to Bregma: A/P: −4.26mm, M/L: +0.6mm, D/V: +4.6mm). Mice were given subcutaneous injections of meloxicam (10 mg/kg) during surgery and 24 & 48 hours after surgery. Mice were group-housed before and after surgery.

**Figure 1:**
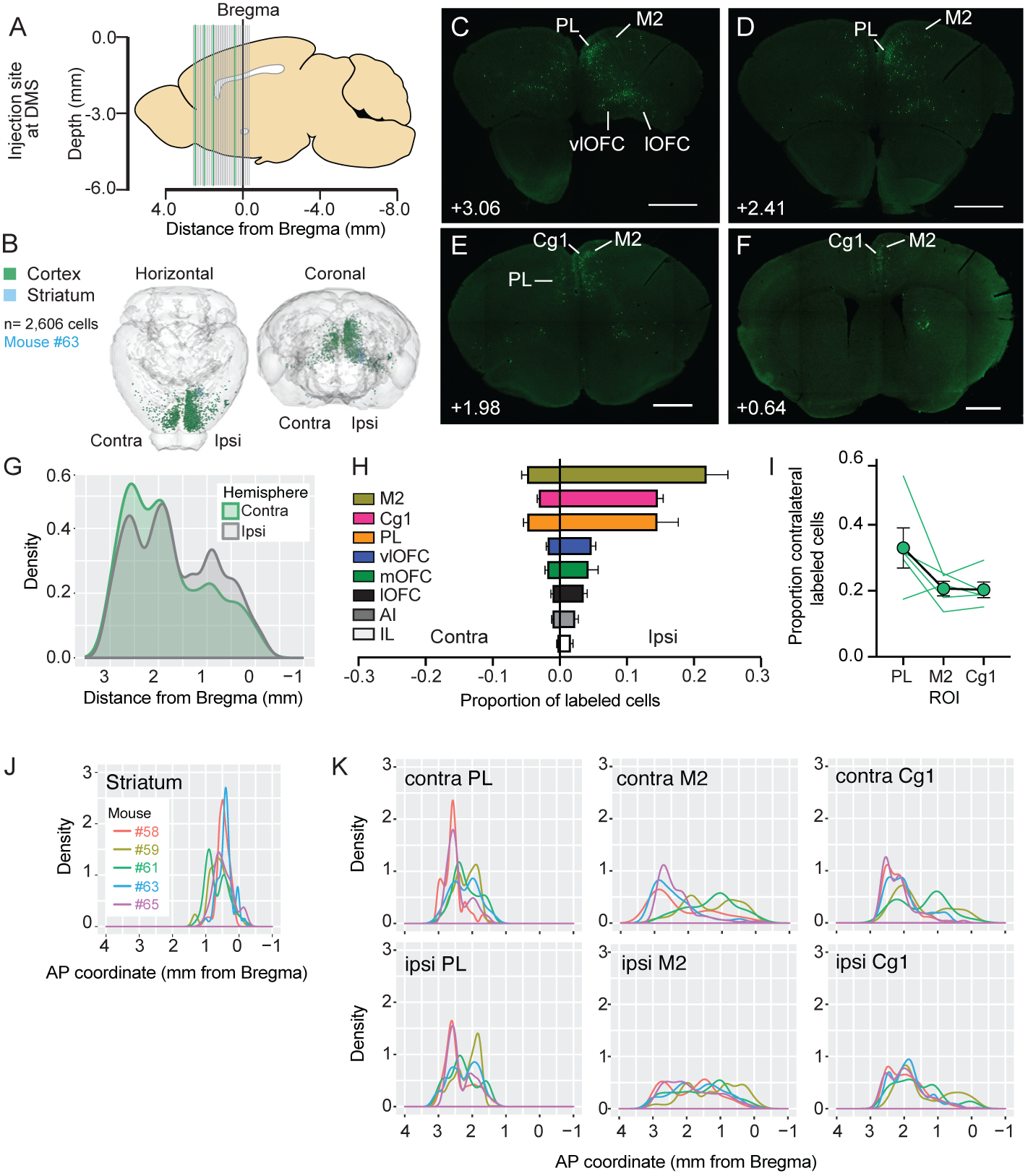
Topography of ipsilateral and contralateral cortical inputs to DMS. (A) Illustration of the anatomical sampling of a subset (n= 4, green vertical lines) of a total of 32 sections (cut thickness: 50 µM; sampling interval: 100 µM) shown in (C-F). (B) 3D visualization of cortical inputs to the DMS in a representative mouse by targeted injection of retro-AAV GFP. Light blue indicates labeled cells within the striatal injection site, while green indicates retrogradely-labeled cortical neurons. (C-F) Representative coronal sections showing labeling of ipsilateral (ipsi) and contralateral (contra) corticostriatal projection neurons to DMS. Abbreviations used are as follows: PL: prelimbic cortex, Cg1: dorsal anterior cingulate cortex, M2: secondary motor area, vlOFC: ventrolateral orbitofrontal cortex, lOFC: lateral orbitofrontal cortex, mOFC: medial orbitofrontal cortex, AI: anterior insula, IL: infralimbic cortex. A/P coordinates relative to Bregma (in mm) are shown for each section. Scale bar, 1000 µM. (G) Probability density plot of anterior to posterior distribution of GFP+ cells by hemisphere (N=5 mice). Fraction of total labeled cells by anatomical region, ipsilateral (ipsi) and contralateral (contra) to the retro-AAV GFP injection site. There was a significant interaction between region and hemisphere (F(1.277, 5.109)=14.66 p= 0.01*; two-way repeated measures ANOVA), indicating that there is regional variation in cSTR projections to contralateral DMS. Fraction of contralateral labeled cells over total by region for PL, M2, and Cg1. There was a trend for the fraction of contralateral labeling to vary by region [F(1.059, 4.236)= 5.84, p= 0.069; one-way repeated measures ANOVA]. (J) Probability density plots by A/P coordinate of labeled cells in DMS injection site by individual mouse. (K) Probability density plots by A/P coordinate of labeled cells in ipsi and contra hemispheres of PL, M2, and Cg1.

**Figure 2:**
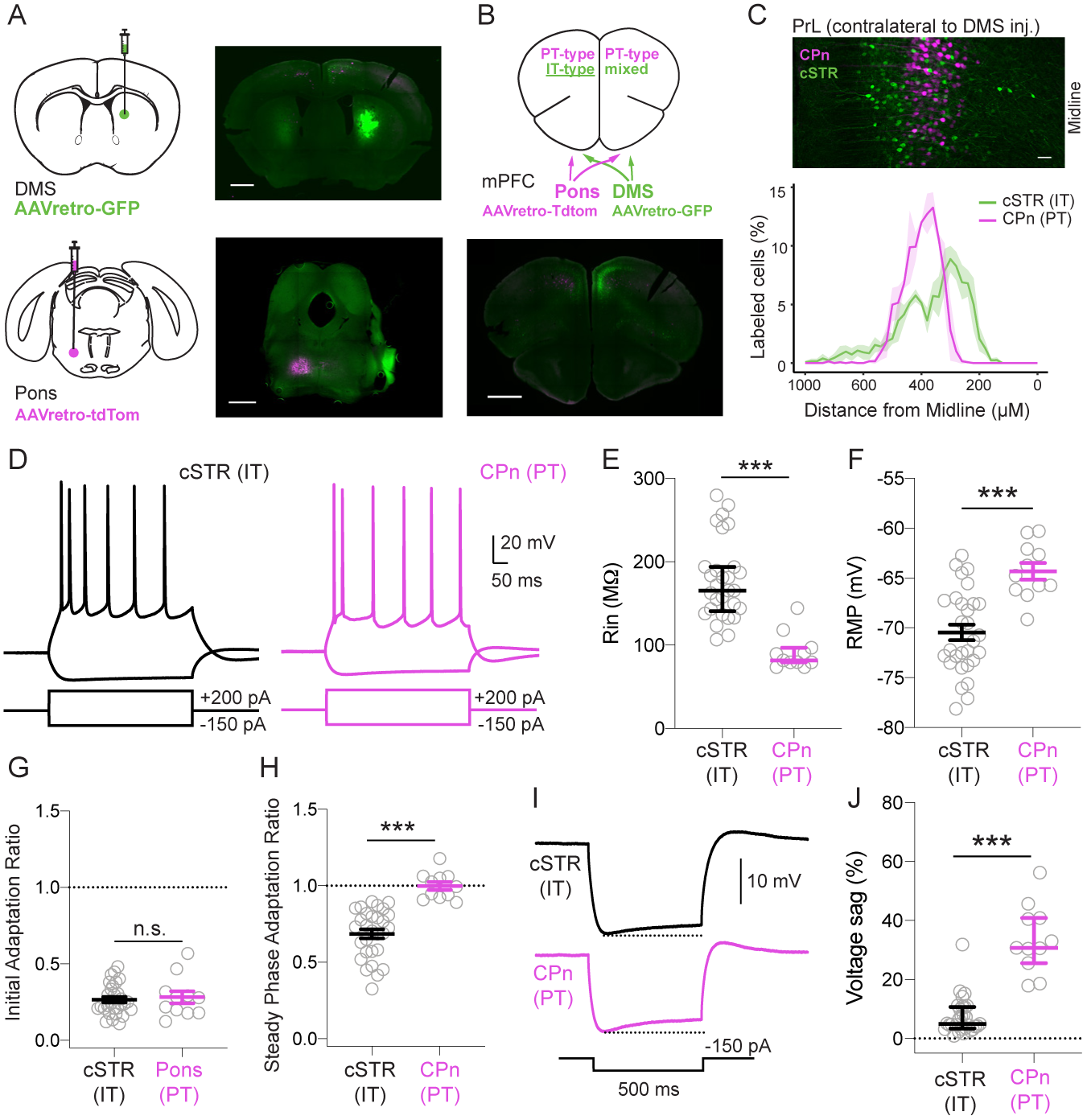
cSTR neurons exhibit properties of IT-type neurons. (A) Schematic of dual infusion of AAVretro-GFP into contralateral DMS and AAVretro-Tdtomato into ipsilateral pons. (B) Viral infusion described in a) are predicted to label non-overlapping populations of cross-corticostriatal (cSTR) IT-type GFP+ neurons and corticopontine (CPn) PT-type Tdtomato+ neurons in contralateral dmP-FC (outlined by white box). (C) Top panel: CPn and cSTR neural populations are non-overlapping in prelimbic (PrL) cortex contralateral to DMS injection. Bottom panel: summary of cSTR and CPn neuron distribution as a function of M/L distance from midline and D/V distance from ventral surface of the brain. Bin size = 25µM. (D) Intrinsic properties and AP firing of cSTR cells (black) and CPn cells (red) in response to depolarizing and hyperpolarizing current steps. (E) CPn cells exhibited significantly lower input resistance (Rin) compared to cSTR neurons. (F) CPn cells were significantly more depolarized at rest compared to cSTR cells. (G) cSTR and CPn cells did not significantly differ in their initial spike adaptation (ISI of initial spike doublet/ last ISI in 5-9 spike train). (H) cSTR cells showed significantly greater spike adaptation during the steady phase of firing compared to CPn cells. (I) Example voltage traces in response to 150 pA hyperpolarizing current shows the presence of H-current mediated voltage sag in CPn but not cSTR cells. (J) Comparison of voltage sag (%) in (I). Data in (E) and (J) presented as median ± interquartile range, otherwise data presented as mean ± SEM. Scale bars in (A,B) = 1000 µM; scale bar in (C) = 50 µM. p<0.001***. Hypothesis tests were conducted using two-tailed student’s t-test (F-H) or Mann-Whitney U test (E, J).

### Gonadectomies

To eliminate gonadal hormone exposure during and after puberty, gonadectomies were performed before puberty onset at P25 as described previously (Delevich et al., 2019c). Briefly, mice were injected with 0.05 mg/kg buprenorphine and 10 mg/kg meloxicam subcutaneously and were anesthetized with 1–2% isoflurane during surgery. The incision area was shaved and scrubbed with ethanol and betadine. Ophthalmic ointment was placed over the eyes to prevent drying. A 1 cm incision was made with a scalpel in the lower abdomen across the midline to access the abdominal cavity. The ovaries or testes were clamped off from the uterine horn or spermatic cord, respectively, with locking forceps and ligated with sterile sutures. After ligation, the gonads were excised with a scalpel. The muscle and skin layers were sutured, and wound clips were placed over the incision for 7–10 days to allow the incision to heal. An additional injection of 10 mg/kg meloxicam was given 12–24 h after surgery. Sham control surgeries were identical to gonadectomies except that the gonads were simply visualized and were not clamped, ligated, or excised. Mice were allowed to recover on a heating pad until ambulatory and were post-surgically monitored for 7–10 days to check for normal weight gain and signs of discomfort/distress. Mice were co-housed with 1-2 siblings who received the same surgical treatment.

### Histology and Fluorescence Microscopy

After eight days of viral expression, mice were transcardially perfused on P29 or P60 for the developmental comparison (Figure 3), or on P60 for the DMS input mapping experiment (Figure 1) and puberty comparison (Figure 4), with 4% paraformaldehyde (PFA) in 0.1M PB (pH=7.4). Brains were harvested and post-fixed in 4% PFA overnight, followed by transfer to a 0.1 M phosphate buffer (PB) solution. For spine imaging experiments, sections were sliced coronally at 150 µM using a vibratome (VT100S Leica Biosystems; Buffalo Grove, IL). For WholeBrain mapping experiments, sections were cut at 50 µM using a vibratome (VT100S Leica Biosystems; Buffalo Grove, IL). Immunohistochemistry was performed for both spine imaging and Wholebrain mapping experiments to enhance the GFP signal (1:1000 chicken anti-GFP, Aves Labs, Inc.; GFP-1020 followed by 1:1000 goat anti chicken AlexaFluor 488, Invitrogen by Thermo Fisher Scientific; A11039). Slides were mounted on slides with Fluoromount-G (Southern Biotech). Coronal sections (A/P: +2.41 mm to +1.40 mm relative to Bregma) were used for spine imaging of the dmPFC and posterior sections of the DMS in the same brain were imaged on an AxioScan Z.1 fluorescent microscope (CRL Molecular Imaging Center, UC Berkeley) to confirm viral targeting.

**Figure 3:**
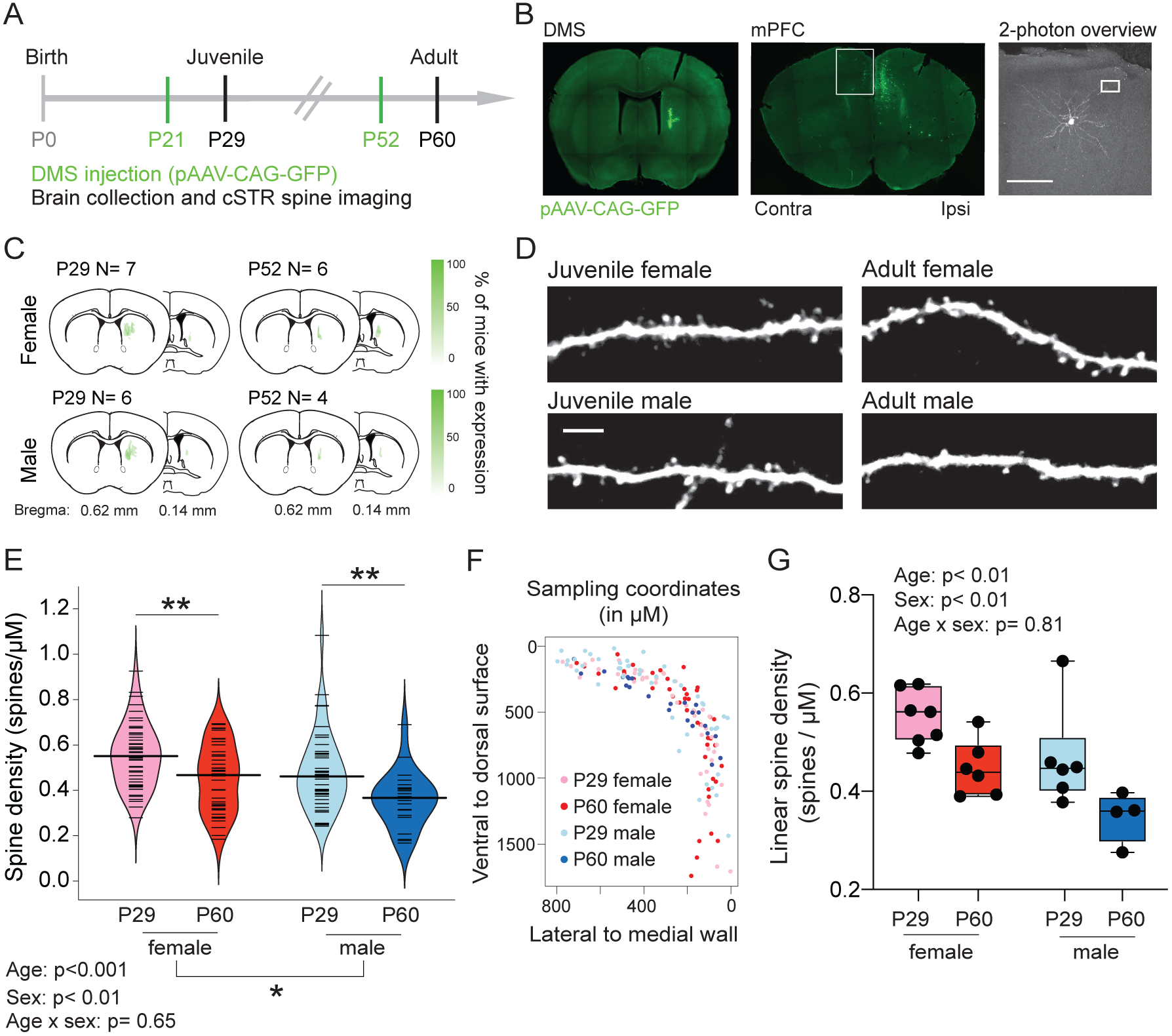
Apical spine density onto cSTR neurons is higher in females compared to males; spine significantly prune across adolescence in both sexes. (A) Stereotaxic cranial injections of pAAV-CAG-GFP were performed on P21 or P52 male and female mice (green). Eight days later at P29 or P60, brains were collected, IHC performed and apical dendrites of cSTR neurons imaged (black). (B) Unilateral infusion of pAAV-CAG-GFP (left panel) led to retrograde expression of GFP in mPFC neurons ipsilateral and contralateral to the injection site (middle panel) (Scale bar = 100 µM). Apical dendrites of GFP+ neurons in contralateral dmPFC (white box; middle panel) were imaged at high magnification on a 2p microscope (white box; right panel). (C) Schematics illustrating center of viral injection site for each mouse; opacity indicates number of mice expressing virus in a given location. (D) Sample images of apical dendrite segments of cSTR neurons in dmPFC (Scale bar = 5 µM). (E) Females exhibited significantly higher apical spine density onto cSTR neurons compared to males, and spine density significantly decreased between P29 and P60 in both sexes. Beanplot shows spine density per cell sampled with median line overlaid. (F) Sampling coordinates for each dendritic segment quantified by group, collapsed across anterior/posterior sections ranging from (+2.41 mm Bregma to +1.40 mm). (G) cSTR spine density significantly differed by age and sex when the linear spine density (total spines/total dendritic length) per animal was compared across groups. **p<0.001, *p<0.01. Hypothesis tests were conducted using linear mixed models in (E) or two-way ANOVA in (G).

**Figure 4:**
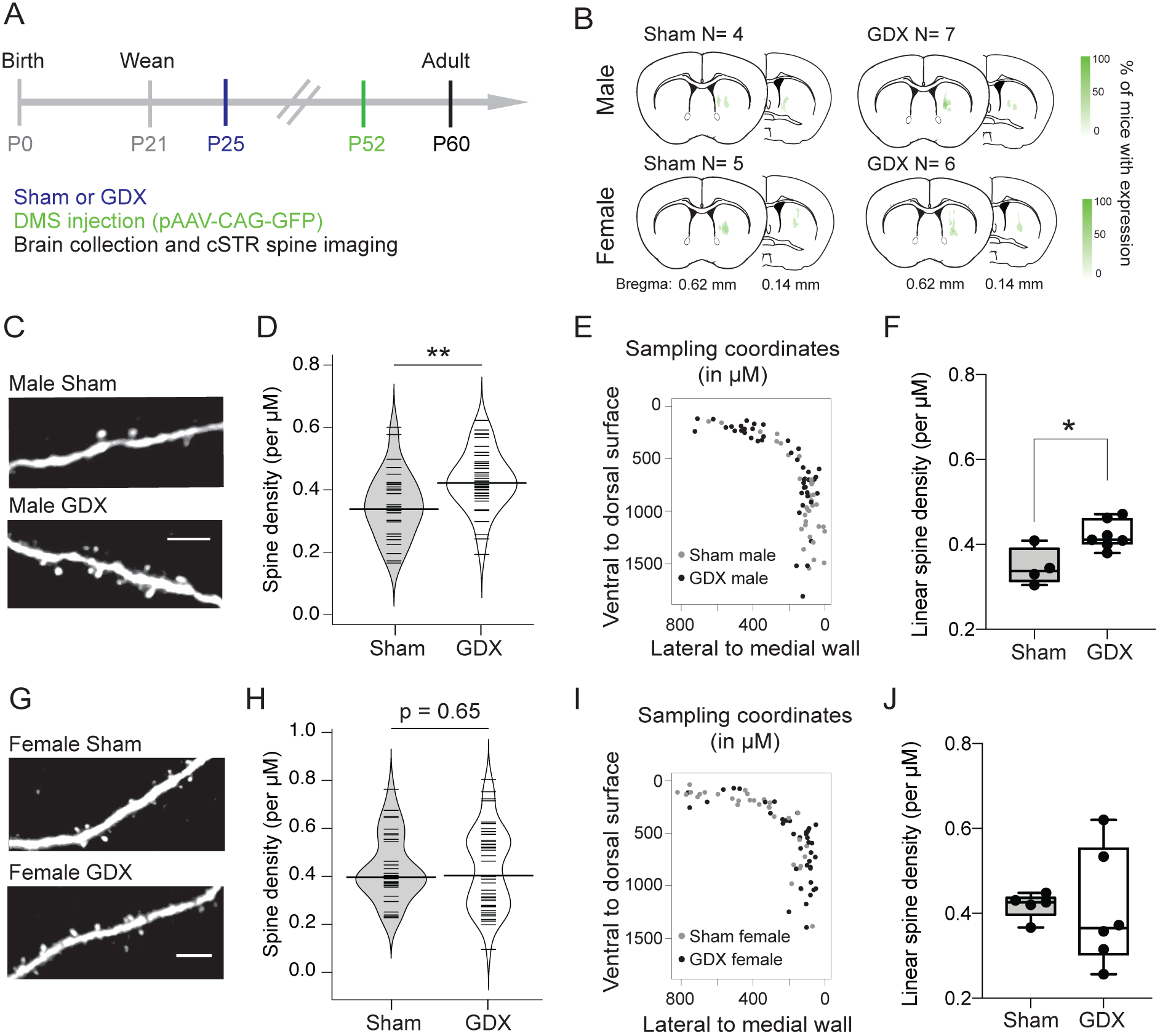
Adult apical spine density onto cSTR neurons is higher in prepubertally GDX males compared to sham males. (A) Sham or GDX surgery was performed in male and female mice at P25 (blue), followed by stereotaxic cranial injections of pAAV-CAG-GFP at P52 (green). Eight days later at P60, brains were collected, IHC performed and apical dendrites of cSTR neurons imaged (black). (B) Schematics illustrating center of viral injection site for each mouse; opacity indicates number of mice expressing virus in a given location. (C) Sample images of apical dendrites on cSTR neurons in male sham (top) and male GDX (bottom) mice. (D) Prepubertal GDX male mice had significantly higher spine density onto cSTR neurons compared to sham males. Beanplot shows spine density per cell sampled with median line overlaid. (E) Sampling coordinates for each dendritic segment quantified by sham and GDX male groups, collapsed across anterior/posterior sections ranging from +2.41 mm Bregma to +1.40 mm. (F) cSTR spine density was significantly higher in GDX males when linear spine density (total spines/total dendritic length) per animal was compared. (G) Sample images of apical dendrites on cSTR neurons in female sham (top) and female GDX (bottom) mice. (H) Spine density onto cSTR neurons did not differ between GDX and sham female mice. Beanplot shows spine density per cell sampled with median line overlaid. (I) Sampling coordinates for each dendritic segment quantified by sham and GDX female groups, collapsed across anterior/posterior sections ranging from +2.41 mm Bregma to +1.40mm. (J) Linear spine density (per animal) did not differ between sham and GDX female groups whereas variance did differ (p<0.05). **p<0.01, *p<0.05. Hypothesis tests were conducted using linear mixed models in (D,H) and two-way ANOVA in (F,J). Scale bar = 5 µM.

For laminar and colocalization analysis of cSTR IT-type neurons and pons-projecting PT-type neurons, we imaged sections on a Zeiss LSM 710 laser scanning confocal microscope using a 10x objective (CNR Biological Imaging Facility, UC Berkeley). For experiments in which the topography of inputs to DMS were mapped (Figure 1), 50 µM coronal sections were processed to enhance GFP signal as described above. Briefly, every other 50 µM section was collected from ∼3.0 to −0.20 mm relative to Bregma, resulting in 100 µM inter-section sampling from anterior frontal cortex through the posterior striatum. A cut was made to identify the injected hemisphere, and sections were mounted and imaged serially at 5x magnification on an AxioScan Z.1 fluorescent microscope (CRL Molecular Imaging Center, UC Berkeley). GFP+ cell bodies were identified and their anatomical position were mapped to Openbrainmap (http://openbrainmap.org) using WholeBrain software (Furth et al., 2018).

For quantitative analysis, the number of input neurons in discrete brain regions was normalized to the total number of input neurons counted. Neurons detected within the striatum were excluded from the total count, as they represented infection within the injection site. In two cases (mouse #58 and #59), there was leak of retro-GFP virus into primary motor cortex that resulted in GFP+ cells in ipsilateral and contralateral primary motor cortex that were absent from cases where no leak occurred. In these cases, we excluded primary motor cortex from the total DMS input counts. Density plots were generated using the geom_density command in the gg-plot2 (Wickham, 2016) package using the default Gaussian smoothing kernel.

### Two photon imaging

Ultima IV laser scanning microscope (Prairie Technologies) and a water immersion 40× magnification 0.8 NA water immersion objective were used to image dendritic spines. A Mai Tai HP laser (Spectra Physics) was tuned to 910 nm for excitation of GFP and 48.301+/-15.739 μM segments of dendrite were imaged at 512 × 512 pixels per inch with resolution 0.0806 μM/pixel and 1 μm z step. In order to quantify apical spines onto cSTR neurons, we imaged apical dendrites within 800 μM from the midline and within 1810 μM of the dorsal pia surface, on the contralateral side relative to injection site. These sampling coordinates encompassed secondary motor cortex (M2), dorsal anterior cingulate cortex (Cg1), and prelimbic cortex (PrL) which together constituted the major input regions to the DMS coordinates we targeted with retro-AAV (Figure 1). Throughout the manuscript we refer to the sampled area collectively as dorsomedial prefrontal cortex (dmPFC).

### Image Processing / Analysis

Dendritic spine analysis was performed blind to the sex / age / surgical treatment of each mouse. Images used for analysis were median-filtered 3-dimensional z stacks. A total of ∼400 µM of dendrite was analyzed for each mouse (see Supplementary table 1). Dendritic spines were scored according to established criteria (Holtmaat et al., 2009) using custom Matlab software (Mathworks, Natick, MA). A dendritic spine was defined by a protrusion greater than 0.4 µM from the dendritic shaft and counts were made according to this definition. The density of dendritic spines, or number of spines per micrometer of dendrite, was recorded. To measure spine density per cell sampled, we recorded the total number of spines over the total length of dendrites per cell sampled. Cell counts comparing the laminar distribution of IT and PT cells were done in IMARIS software (IMARIS). Counts were performed using 3D images that were obtained with Z stack function on confocal microscope.

**Table 1:** Electrophysiological properties of cSTR neurons.

|  | Juvenile Female<br>(n= 10) | Juvenile Male<br>(n=11) | Adult Female<br>(n=15) | Adult Male<br>(n=14) | Statistics |
| --- | --- | --- | --- | --- | --- |
| RMP (mV) | -67.86 ± 1.50 | -70.63 ± 0.98 | -70.95 ± 1.09 | -69.94 ± 1.13 | p>0.05 |
| Rin (MΩ) | 177.8 ± 15.31 | 193.7 ± 15.38 | 163.5 ± 8.24 | 192.3 ± 15.45 | p>0.05 |
| Max frequency (Hz) | 30.6 ± 0.95 | 28.4 ± 1.72 | 29.9 ± 1.4 | 28.6 ± 1.8 | p>0.05 |
| Initial adaptation | 0.26 ± 0.02 | 0.34 ± 0.04 | 0.24 ± 0.03 | 0.29 ± 0.02 | **Sex: \*p<0.05** |
| Steady phase adaptation | 0.83 ± 0.03 | 0.81 ± 0.03 | 0.66 ± 0.05 | 0.72 ± 0.03 | **Age: \*\*p<0.01** |
| Sag (%) | 0.51 ± 0.11 | 0.64 ± 0.13 | 0.76 ± 0.19 | 0.74 ± 0.13 | p>0.05 |

For image presentation (Figure 3D, Figure 4C, G) the relevant sections of dendrite were projected onto a 2-dimensional image, which was then Gaussian filtered and contrasted for presentation. To project the relevant sections of dendrite onto a 2-dimensional image, the frames from the 3-dimensional z stack in which spines were most clearly in focus were combined using the maximum intensity z projection functions in ImageJ.

### Slice electrophysiology

Early adolescent (P29-P31) and adult mice (P60-90) were deeply anesthetized with an overdose of ketamine/xylazine solution and perfused transcardially with ice-cold cutting solution containing (in mM): 110 choline-Cl, 2.5 KCl, 7 MgCl2, 0.5 CaCl2, 25 NaHCO3, 11.6 Na-ascorbate, 3 Na-pyruvate, 1.25 NaH2PO4, and 25 D-glucose, and bubbled in 95% O2/ 5% CO2. 300 µm thick coronal sections were cut in ice-cold cutting solution before being transferred to ACSF containing (in mM): 120 NaCl, 2.5 KCl, 1.3 MgCl2, 2.5 CaCl2, 26.2 NaHCO3, 1 NaH2PO4 and 11 Glucose. Slices were bubbled with 95% O2/ 5% CO2 in a 37°C bath for 30 min, and allowed to recover for 30 min at room temperature before recording at 32°C.

Whole-cell recordings were obtained from visually identified IT-cSTR neurons across all layers of the prelimbic and dorsal anterior cingulate subdivision of mPFC. IT-cSTR neuronal identity was established by the presence of GFP in the hemisphere contralateral to the GFP retro-AAV injection site in DMS. Whole-cell current clamp recordings were performed using a potassium gluconate-based intracellular solution (in mM): 140 K Gluconate, 5 KCl, 10 HEPES, 0.2 EGTA, 2 MgCl2, 4 MgATP, 0.3 Na2GTP, and 10 Na2-Phosphocreatine. All recordings were made using a Multiclamp 700 B amplifier and were not corrected for liquid junction potential. Data were digitized at 20 kHz and filtered at 2 kHz using a Digidata 1440 A system with pCLAMP 10.2 software (Molecular Devices, Sunnyvale, CA, USA). Only cells with series resistance of < 25 MΩ were retained for analysis, and series resistance was not compensated. Cells were discarded if parameters changed more than 20%. Data were analyzed using pClamp software.

Current clamp recordings were performed in the presence of NBQX (10 µM), AP5 (50 µM), and Picrotoxin (10 µM). Briefly, neurons were injected with 500 ms current steps starting from −200 pA with increasing 50 pA steps until a maximum of 500 pA. Input resistance was measured using the steady-state response to a 100 pA hyperpolarizing current step. The membrane time constant (tau) was obtained from exponential fits to the −50 pA current step. Voltage sag was measured as the percent change from the peak potential to the steady state potential during a 150 pA negative current injection step. The adaptation index was calculated as the ratio of the second interspike interval (ISI) and the final ISI in response to a 500 ms, 450 pA current injection. Data were collected from at least 2 slices per animal.

### Statistics

Because spine density measures included multiple cells from the same brain, these data were analyzed by fitting a linear mixed model using residual maximum likelihood. The “nmle” package was used for linear mixed effects modeling and “beanplot” package was used to create density plots of our data (RStudio). Transformed values (square root of the raw spine densities) were used when the residuals of models that used untransformed data significantly deviated from normality. Sex and age (developmental comparison) or surgical procedure (puberty comparison) were held as the fixed effects and the animal sampled was held as the random effect. Normality of the residuals for each model was confirmed using the Shapiro-Wilks test. Anova was performed to distinguish the model that best explained the variance in our data. Statistical analyses were performed using R Studio. Mean ± SEM was reported for each group, unless otherwise noted. Statistical significance is marked as * for p < 0.05, ** for p <0.01, and *** for p<0.001.

## RESULTS

### Topography of corticostriatal projections to DMS

In order to selectively image IT-type neurons in the dorsal medial prefrontal cortex (dmPFC), we targeted retro-AAV to the dorsomedial striatum (DMS). Unilateral infusion of retro-AAV into DMS resulted in labeled cells in the ipsilateral and contralateral hemispheres, with cross-corticostriatal neurons (cSTR) in the contralateral hemisphere ostensibly representing a uniquely IT-type population (Wilson, 1987). We first examined the topography of neurons labeled by retro-AAV-GFP infusion into DMS. Serial 50 µM coronal sections were prepared and imaged on an Axio ScanZ.1 microscope.

GFP+ neurons in the rostral portion of the brain (approximately +3.0 to −0.20 mm relative to Bregma) were quantified and registered to a reference atlas using the WholeBrain software package (Furth et al., 2018) (Figure 1A,B). We observed that the topography of cSTR neurons was shifted rostrally compared to ipsilateral corticostriatal projection neurons (Figure 1B-G). When we analyzed the fraction of GFP+ cells by anatomical region, we found that there was a significant interaction between region and hemisphere (F(1.277, 5.109)=14.66 p= 0.01, two-way repeated measures ANOVA), suggesting that there is regional variation in cSTR projections to DMS (Figure 1H). Together, prelimbic (PL), dorsal anterior cingulate (Cg1), and secondary motor cortex (M2) contributed 64 ± 2% (mean ± SEM; N=5) of the total input to DMS (Figure 1H). Among these top three input regions, we analyzed the fraction of GFP+ cells located in the contralateral hemisphere and observed a trend-level effect of ROI (F(1.059, 4.236)= 5.84, p= 0.069; one-way repeated measures ANOVA) (Figure 1I). Finally, we compared the location of GFP+ cells in striatum, representing the injection site (Figure 1J) and separately plotted the A/P topographical distribution of GFP+ cells in the ipsilateral and contralateral hemispheres by regions which revealed unique topographies, with more rostral peaks for the contralateral hemisphere (Figure 1K).

### DMS-targeted retro-AAV labels IT-type pyramidal neurons in contralateral dmPFC

To confirm that unilateral retro-AAV infusion to the DMS selectively labels IT-type neurons in contralateral dmPFC, we injected retro-AAV-GFP into the left striatum and retro-AAV-tdTomato into the right pons of the same mice (Figure 2A). Based on anatomical projections, we predicted that GFP+ neurons in the left dmPFC are a mix of PT-type and IT-type neurons (Levesque et al., 1996; Kita and Kita, 2012) whereas GFP+ neurons in the right dmPFC should be purely IT-type (Figure 2B). Meanwhile, Tdtomato+ neurons in both hemispheres are expected to be PT-type.

We found that within left dmPFC Tdtomato+ corticopontine (CPn) neurons did not overlap with GFP+ cross-corticostriatal (cSTR) projection neurons (Figure 2C). Furthermore, cSTR and CPn neurons exhibited distinct laminar distributions in the prelimbic cortex (PL) (Figure 2C). GFP+ cSTR neurons localized to both more superficial and deep coordinates compared to Tdtomato+ CPn neurons, but these two populations significantly overlapped 300-550 µM from the midline (Figure 2C), consistent with previous studies (Morishima and Kawaguchi, 2006; Anderson et al., 2010; Dembrow et al., 2010; Kiritani et al., 2012; Li et al., 2015). Finally, in a separate experiment, we co-injected PL and DMS and observed that a subset of neurons in contralateral PL were co-labeled (Figure S1), consistent with data showing that single corticostriatal neurons project to both contralateral cortex and contralateral striatum (Winnubst et al., 2019).

Cortical IT-type and PT-type neurons exhibit characteristic intrinsic electrophysiological properties (Hattox and Nelson, 2007; Dembrow et al., 2010; Oswald et al., 2013). We performed whole-cell current clamp recordings on GFP+ cells in dmPFC of adult (∼P60) mice injected with retroAAV-GFP into ipsilateral Pons (CPn) or contralateral DMS (cSTR) in the presence of the synaptic blockers AP5 (50 µM), NBQX (10 µM), and PTX (10 µM) (Figure 2D). Consistent with previous reports, CPn neurons had significantly lower input resistance compared to cSTR neurons (cSTR 165.5 ± 53.2 MΩ vs. Pons 73.52 ± 17.53 MΩ; U= 11, p<0.0001***) (Figure 2E). In addition, CPn neurons were significantly more depolarized at rest compared to cSTR neurons (cSTR −70.46 ± 0.77 mV vs. Pons −64.32 ± 0.83 mV; t_38_= 4.5, p<0.0001***) (Figure 2F).

We observed that both cSTR and CPn neurons exhibited doublet spikes at the onset of the 500 ms square current injection (Figure 2G), and there was no significant difference between cSTR and CPn in the ratio between the first interspike interval (ISI) and last ISI, referred to as the initial adaptation ratio (cSTR 0.27 ± 0.019 vs. Pons 0.28 ± 0.040; t38= 4.15, p= 0.68) (Figure 2G). However, only cSTR exhibited spike adaptation during the steady phase of spiking (cSTR 0.69 ± 0.30 vs. Pons 1.00 ± 0.03; t_38_= 6.12, p<0.0001***) (Figure 2H). Finally, PT-type neurons are distinguished by prominent voltage sag in response to hyperpolarizing current steps that indicate the presence of hyperpolarization-activated currents (I_h_) (Figure 2I). CPn neurons showed significantly greater voltage sag (%) in response to a 150 pA hyperpolarizing current step compared to cSTR-projecting neurons (cSTR 4.94 ± 7.3 vs. CPn 30.7 ± 15.3; U= 6, p<0.0001***) (Figure 2J). Together the intrinsic electrophysiological properties of cSTR neurons labeled by retro-AAV-GFP were consistent with IT-type cortical projection neurons.

### Apical dendritic spines onto cSTR neurons significantly prune across adolescence and differ by sex

To determine whether apical spine density onto cSTR neurons changes during adolescence, we injected retro-AAV-GFP into the left DMS and used two-photon microscopy to image dendrites in the dmPFC within the right hemisphere (Figure 3B). Groups included female (n=7) and male (n=6) juvenile (P29) mice and female (n=6) and male (n=4) early adult (P60) mice. Our sampled region of dmPFC encompassed prelimbic cortex (PL), dorsal anterior cingulate cortex (Cg1), and secondary motor cortex (M2) (see methods for more details) (Figure 3F). All four groups were comparably sampled in the region of interest (Figure 3F). We hypothesized that IT cSTR spine density would decrease from P29 to P60, as has been shown in PT neurons across adolescence in both male (Johnson et al. 2016) and female (Boivin et al. 2018) mice. We observed that P60 adult mice exhibited lower spine density, (0.4183 ± 0.0181 spines / µM) compared to P29 juvenile mice, (0.5171 ± 0.0161 spines / µM) (t_20_= −4.291, p= 0.0004**; Figure 3E). In addition, we found that P29 females (0.5503 ± 0.0195 spines / µM) exhibited higher spine densities than P29 males (0.4795 ± 0.0252 spines / µM). This sex difference persisted into early adulthood, as P60 females (0.4522 ± 0.0232 spines / µM) exhibited higher spine densities compared to P60 males (0.3593 ± 0.0250 spines / µM) (t_20_=−3.262, p= 0.0039*; Figure 3E). Meanwhile, there was no significant interaction between age and sex (t_19_=−0.466, p =0.647), suggesting that the extent to which spines are pruned is comparable between sexes. These results were consistent when we compared linear spine density measures by animal (main effect of sex: t_20_=−3.118, p=0.0054**; main effect of age: t_20_=−3.819, p=0.0011**; no interaction between age and sex: t_19_=−0.245, p=0.809) (Figure 3G).

### cSTR neurons exhibit subtle age and sex differences in spike frequency adaptation

We next performed whole-cell current clamp recordings onto GFP+ cSTR neurons in the dmPFC of juvenile (P29-31) and adult (P60-62) male and female mice to determine whether intrinsic properties differed by sex or age. We found that with the exception of spike frequency adaptation, cSTR intrinsic properties did not differ by age or sex (Table 1). We found a significant effect of sex on initial adaptation index, with females exhibiting shorter interspike interval (ISI) of initial doublet spikes compared to males (F(1, 46)= 5.33, p= 0.026*) (Table 1). Next, we found that there was a significant effect of age on the steady phase adaptation index, with adults exhibiting significantly greater steady phase adaptation compared to juveniles (F(1, 46)= 11.06, p= 0.0017**). (Table 1). Spike frequency adaptation is attributed to K+ currents activated by either calcium (IAHP) or voltage-activated K+ (IM) currents (Ha and Cheong, 2017). Finally, there was a significant interaction between sex and current on firing rate, suggesting that females exhibit a greater gain in firing rate at large current steps compared to males (Figure S2).

### Prepubertally gonadectomized males exhibit higher spine densities in early adulthood compared to gonadally intact controls

To determine whether pubertal development plays a role in spine pruning across adolescence, we compared spine densities onto cSTR neurons in adult mice that were gonadectomized before puberty (P24-25) (7 males, 6 females) or received sham surgery (4 males, 5 females) (Figure 4A,B, Figure S3). We found that pre-pubertally gonadectomized males had a significantly higher spine density (0.6515 ± 0.0110 spines / µM) onto dmPFC cSTR neurons compared to age-matched sham controls (0.5852 ± 0.0150 spines / µM) at P60 (t_9_ = 3.650, p= 0.0053**) (Figure 4D). This effect was also significant when the total linear spine density per animal was compared across groups (t_9_ =−3.206, p= 0.011*) (Figure 4F).

Spine density onto cSTR neurons in prepubertally gonadectomized females (0.6459 ± 0.0204 spines / µM) did not significantly differ from sham controls (0.6507 ± 0.0181 spines / µM) at P60 (t_9_ =−0.4645, p= 0.6533) either when density measures were compared by cells (Figure 4H) or animal (t_9_ = −0.137, p= 0.894) (Figure 4J). However, the variance of GDX female spine density was significantly higher compared to that of sham females (F(5, 4)= 20.28, p= 0.012*) (Figure 4J). To more directly examine how gonadectomy affected adolescent spine pruning, we normalized spine density measures to juvenile measures and compared intact and GDX groups within the same sex (Figure 5A, B). Unmanipulated adult and sham adult data were pooled because they did not significantly differ (Supplementary Figure 4). We found that both juvenile male and GDX male spine density were significantly different from intact adult male spine density, whereas GDX males did not significantly differ from juvenile males (Figure 5A). Meanwhile, both intact and GDX female spine density significantly differed from juvenile female spine density (Figure 5B).

**Figure 5:**
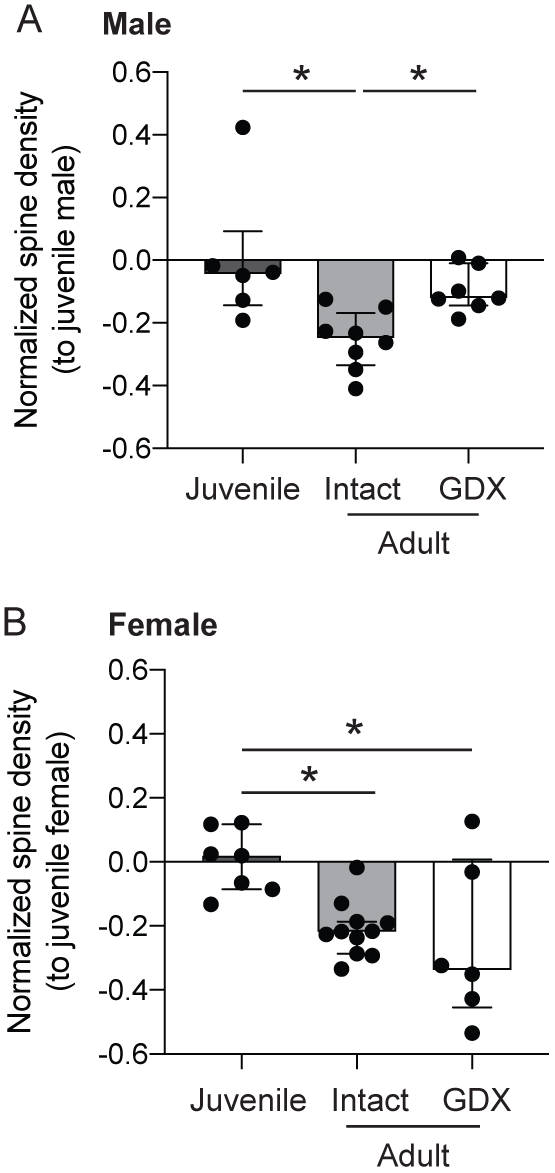
Comparison to juvenile apical spine density data suggests that GDX males prune significantly less during adolescence compared to intact males. (A) Apical spine density values onto cSTR neurons normalized to juvenile male values. (B) Apical spine density values onto cSTR neurons normalized to juvenile female values. Data presented as median ± IQR. Hypothesis tests were conducted using Kruskal-Wallis one way ANOVA with post hoc Dunn’s test for multiple comparisons. *p<0.05.

## DISCUSSION

Our current study focused on the adolescent maturation of a specific cortical pyramidal cell type that projects to the dorsomedial striatum via the corpus callosum (referred to as cross-corticostriatal or cSTR neurons). These cSTR neurons represent a subpopulation of the intratelencephalic (IT-type) cortical projection neurons. First, we characterized the topography of corticostriatal projections to the DMS. We found evidence that cSTR neurons are distributed more rostrally compared to ipsilateral corticostriatal projection neurons. In addition, we found evidence that the relative contribution of ipsilateral vs. contralateral inputs significantly varied by region. Consistent with previous studies, we found that the major input regions to the DMS included M2, Cg1, and PL (Gabbott et al., 2005; Pan et al., 2010; Hintiryan et al., 2016; Hunnicutt et al., 2016) which we referred to as dmPFC here – for discussion of this nomenclature, refer to (Laubach et al., 2018).

When we analyzed the relative contribution of cSTR projections by region, we observed a trend for enriched cSTR inputs from the PL compared to M2 and Cg1. Next, we confirmed that cSTR neurons labeled via retrograde AAV injection into contralateral DMS exhibit properties consistent with IT-type neurons, including non-overlap with PT-type neurons and electrophysiological properties such as spike frequency adaptation and low voltage-sag. We then focused on spine density on the apical tufts of dmPFC cSTR neurons in pre and postpubertal mice (ages P29 and P60). We found that males at both ages exhibited significantly lower spine density compared to females, but both sexes pruned to a similar extent between P29 and P60. Finally, we found that prepubertal gonadectomy in males was associated with significantly higher spine density in adulthood, suggesting that pubertal development plays a causal role in spine pruning onto these neurons in males. In females, no significant difference in spine density was found when GDX and sham adults were compared, although the variance of spine density was significantly higher in GDX females compared to sham.

Our last finding raises the question: by which mechanism does pubertal development promote dendritic spine pruning in males? The localization of steroid hormone receptors in the frontal cortex provide insight into potential mechanisms by which testicular hormones may act. Importantly, the brains of male and female mice express steroid hormone receptors for both androgens and estrogens (Piekarski et al., 2017b), and testosterone can be converted to estradiol via aromatization (Nelson and Bulun, 2001; Shay et al., 2018). Furthermore, the aromatase enzyme is present in the frontal cortex (Akther et al., 2015; Mitra et al., 2015). In the frontal cortex of rodents, estrogen receptor beta (ERbeta) receptors primarily localize to parvalbumin expressing fast-spiking interneurons in males and females (Blurton-Jones and Tuszynski, 2002; Kritzer, 2002). In female rodents, it has previously been shown that ovarian hormones can modulate inhibitory synaptic transmission (Piekarski et al., 2017a) and, more specifically, the intrinsic excitability of fast-spiking parvalbumin interneurons (Clemens et al., 2019; Delevich et al., 2019a). Steroid hormone receptors are also expressed by microglia (Sierra et al., 2008), whose role in synaptic pruning has been described (Paolicelli et al., 2011). Interestingly, a recent paper demonstrated that microglia engulfment of dendritic spines in L5 of PL significantly increased between P24 and P39, roughly coinciding with pre- and post-pubertal time points in rats (Mallya et al., 2019), although the authors did not assess pubertal status in this study. If and how gonadal hormones regulate the function of microglia during puberty is an exciting area for future study. Alternatively, it is possible that differences in spine density between sham and GDX males occur downstream of behavioral (Jardi et al., 2018; Delevich et al., 2019c) or metabolic consequences (Gentry and Wade, 1976; Krotkiewski et al., 1980; Inoue et al., 2010) of prepubertal gonadectomy.

In females, we did not observe a significant difference between sham and GDX mice in adulthood, but GDX was associated with significantly higher within-group variance in apical dendritic spine density. This makes these data more difficult to interpret. The spine density average suggests that ovarian hormones do not play a significant role in spine pruning on the cSTR cell type, however, the bimodal distribution and higher variance raise the question of whether an underlying factor stemming from gonadal hormone manipulations drove new variability. Therefore, in addition to the null hypothesis we considered two speculative explanations. First, it is possible that GDX at P25 did not occur early enough to prevent earliest pubertal ovarian hormone secretion in all mice. Second, it is possible that GDX in females affected alternative mechanism of hormone production which in turn affected spine density. Thus, given the significant change in variance, we cannot fully rule out a role for pubertal processes in regulating spine density on cSTR neurons in females. Indeed, a previous study found that ovariectomy in adulthood was associated with reduced apical spine density onto L2/3 pyramidal neurons in the mPFC of rats (Wallace et al., 2006). Future experiments in females should further explore the influence of various sources of steroid sex hormones at different developmental periods (i.e. pre- and post-pubertal) on cell-type specific spine pruning.

While we focused on cSTR neurons, we and others have shown that cSTR neurons overlap with cortico-cortical IT-type projection neurons (Sohur et al., 2014; Winnubst et al., 2019), suggesting that our results may apply more broadly to IT-type neurons within mPFC or broader frontal cortices. On the other hand, even within mPFC significant heterogeneity has been reported among IT-type neurons (Morishima and Kawaguchi, 2006; Anastasiades et al., 2019). Consistent with previous studies, we found that dual infusion of retro-AAV into the pons and contralateral striatum labeled two intermingled but non-overlapping populations of pyramidal neurons in layer 5 of PL. Cross-corticostriatal (cSTR) and corticopontine (CPn) neurons exhibited the characteristic intrinsic properties of IT-type and PT-type neurons, respectively. We observed that in mouse, both cell types exhibited initial doublet spikes, whereas a previous report observed this only in CPn neurons in rats, highlighting a potential species difference (Morishima and Kawaguchi, 2006). We observed that the topography of cSTR neurons was shifted more rostrally compared to ipsilateral corticostriatal projection neurons. Interestingly, a previous study using anterograde tracing found that frontal cortical regions (including M2) exhibited more cSTR projections compared to sensory and motor regions (Hooks et al., 2018). Furthermore, compared to more caudal motor and sensory areas, a greater proportion of M2 corticostriatal projections originate from IT-type neurons relative to PT-type neurons (Hooks et al., 2018).

These data raise questions regarding the function of cSTR neurons and the significance of their enrichment in frontal cortical regions. At the microcircuit level, connectivity between IT-type corticostriatal neurons and PT-type neurons is highly asymmetric, with IT-type neurons exerting greater influence over PT-type neurons (Morishima and Kawaguchi, 2006; Brown and Hestrin, 2009; Kiritani et al., 2012). Furthermore, studies in motor cortices have shown that IT-type neural activity is associated with planning/motor preparation and corticospinal activity with motor execution (Bauswein et al., 1989; Turner and DeLong, 2000; Li et al., 2015), suggesting that ‘output-potent’ PT-type activity may be inherited from preparatory IT-type activity (Kaufman et al., 2014; Li et al., 2015). Therefore, adolescence-associated changes in spine density onto cSTR neurons could influence how inputs to the mPFC are integrated and modulate downstream targets, including local PT-type neurons and striatal populations.

Across the adolescent transition, there is evidence that behavior becomes less stimulus-directed and more goal-directed (Sturman et al., 2010; Andrzejewski et al., 2011; Blakemore and Robbins, 2012; Naneix et al., 2012; Hammerslag and Gulley, 2014). Here, we show that DMS-projecting cSTR neurons, which are implicated in goal-directed learning (Hart et al., 2018a; Hart et al., 2018b), exhibit significant sex differences in apical dendritic spine density. Furthermore, we show that these cSTR neurons prune in both sexes across adolescence and in males this loss of spines is dependent on the presence of the gonads at puberty. Our study adds to a growing body of research demonstrating that spine density and dynamics change within agranular frontal cortices during adolescence (Zuo et al., 2005; Gourley et al., 2012; Johnson et al., 2016; Boivin et al., 2018), and provides novel evidence that spine pruning onto a projection-defined cell-type in the frontal cortex is sensitive to pubertal development. More work needs to be done in order to understand how spine pruning relates to changes in neural computation and cognitive processes during adolescence (Selemon, 2013), but recent studies have made progress, associating changes in spine density onto OFC neurons to alterations in goal-directed behavior (DePoy et al., 2019; Hinton et al., 2019). Data in humans suggests that pubertal status and circulating gonadal hormone levels influence working memory use during learning in both sexes (Master et al., 2019), sensitivity to immediate rewards during intertemporal decision-making in males (Laube et al., 2017), and a variety of social and affective behaviors in both sexes (Vijayakumar et al., 2018; Goddings et al., 2019). Future work will be needed to test whether developmental changes in cSTR structure and function relate to developmental changes in goal-directed learning or other behaviors.

At a more macroscopic level, our study highlights a cell-type that may be important for understanding sex differences in frontal cortex function and disease risk, particularly psychiatric diseases that emerge during adolescence (Paus et al., 2008). Importantly, humans possess IT-type and PT-type glutamatergic neurons that are homologous to the cell-types studied here (Lake et al., 2016; Hodge et al., 2019), although it has been argued that PT-type neurons are more abundant in mice compared to humans (Hodge et al., 2019). Regardless, the evolutionary conservation of cortical cell types may enable researchers to infer properties of human neuronal cell types and make predictions about their developmental trajectories. Translational research will be critical to our understanding of how puberty influences brain maturation in humans, where investigation into the effects of gonadal hormone levels are, by necessity, correlational. Several observations suggest that IT-type neurons may exhibit a more protracted maturation compared to PT-type neurons. Postmortem data from nonhuman primates and humans suggest that cortico-cortical projection neurons located in superficial cortical layers may undergo more marked pruning of supernumerary synapses over the course of postnatal development (Bourgeois et al., 1994). Data from rats show that considerable dendritic ramification of layer 2/3 mPFC neurons occurs during adolescence, taking place earlier in females compared to males, potentially reflecting differences in pubertal timing (Markham et al., 2013). In addition, slice electrophysiology experiments in rodents and nonhuman primates suggest that synaptic inhibition onto IT-type neurons in frontal regions matures during the peripubertal period (Gonzalez-Burgos et al., 2015; Vandenberg et al., 2015; Piekarski et al., 2017a), which could in turn impact dendritic spine pruning (Chen et al., 2018; Ng et al., 2018).

Here, we add to this growing body of evidence by showing that apical spines onto cSTR neurons prune during adolescence and providing evidence that this process may be regulated by puberty in males. Protracted, pubertal-sensitive maturation of cSTR neurons may have important implications for prefrontal cortex function both in relation to the adaptive emergence of adult-like behavior and the maladaptive susceptibility to psychiatric disorders that emerge post-puberty.

## ACKNOWLEDGEMENTS

We thank Dr. Daniel Furth for generous technical assistance with Wholebrain analysis. We thank Dr. David Piekarski and Dr. Lance Kriegsfeld for training and assistance with gonadectomies and the Wilbrecht lab for discussion. Confocal imaging experiments were conducted at the CNR Biological Imaging Facility. Research Slidescanner imaging experiments were conducted at the CRL Molecular Imaging Center, supported by the Biological Faculty Research Fund. We would like to thank Holly Aaron and Feather Ives for their microscopy training and assistance. This project was supported by a seed grant from the University of California Developmental Science of Adolescence Consortium.

This preprint was formatted using a template created by David Paquet (available:https://github.com/drpaquet/biorxiv_preprint_templates)

## REFERENCES

Akther S, Huang Z, Liang M, Zhong J, Fakhrul AA, Yuhi T, Lopatina O, Salmina AB, Yokoyama S, Higashida C, Tsuji T, Matsuo M, Higashida H (2015) Paternal Retrieval Behavior Regulated by Brain Estrogen Synthetase (Aromatase) in Mouse Sires that Engage in Communicative Interactions with Pairmates. Front Neurosci 9:450.

Anastasiades PG, Boada C, Carter AG (2019) Cell-Type-Specific D1 Dopamine Receptor Modulation of Projection Neurons and Interneurons in the Prefrontal Cortex. Cereb Cortex 29:3224–3242.

Anderson CT, Sheets PL, Kiritani T, Shepherd GM (2010) Sublayer-specific microcircuits of corticospinal and corticostriatal neurons in motor cortex. Nat Neurosci 13:739–744.

Anderson SA, Classey JD, Conde F, Lund JS, Lewis DA (1995) Synchronous development of pyramidal neuron dendritic spines and parvalbumin-immunoreactive chandelier neuron axon terminals in layer III of monkey prefrontal cortex. Neuroscience 67:7–22.

Andrzejewski ME, Schochet TL, Feit EC, Harris R, Mckee BL, Kelley AE (2011) A Comparison of Adult and Adolescent Rat Behavior in Operant Learning, Extinction, and Behavioral Inhibition Paradigms. Behavioral Neuroscience 125:93–105.

Baker A, Kalmbach B, Morishima M, Kim J, Juavinett A, Li N, Dembrow N (2018) Specialized Subpopulations of Deep-Layer Pyramidal Neurons in the Neocortex: Bridging Cellular Properties to Functional Consequences. J Neurosci 38:5441–5455.

Bauswein E, Fromm C, Preuss A (1989) Corticostriatal cells in comparison with pyramidal tract neurons: contrasting properties in the behaving monkey. Brain Res 493:198–203.

Blakemore SJ, Robbins TW (2012) Decision-making in the adolescent brain. Nature Neuroscience 15:1184–1191.

Blurton-Jones M, Tuszynski MH (2002) Estrogen recep-tor-beta colocalizes extensively with parvalbumin-labeled inhibitory neurons in the cortex, amygdala, basal forebrain, and hippocampal formation of intact and ovariectomized adult rats. J Comp Neurol 452:276–287.

Boivin JR, Piekarski DJ, Thomas AW, Wilbrecht L (2018) Adolescent pruning and stabilization of dendritic spines on cortical layer 5 pyramidal neurons do not depend on gonadal hormones. Dev Cogn Neurosci 30:100–107.

Bourgeois JP, Goldman-Rakic PS, Rakic P (1994) Synaptogenesis in the prefrontal cortex of rhesus monkeys. Cereb Cortex 4:78–96.

Bramen JE, Hranilovich JA, Dahl RE, Forbes EE, Chen J, Toga AW, Dinov ID, Worthman CM, Sowell ER (2011) Puberty influences medial temporal lobe and cortical gray matter maturation differently in boys than girls matched for sexual maturity. Cereb Cortex 21:636–646.

Brown SP, Hestrin S (2009) Intracortical circuits of pyramidal neurons reflect their long-range axonal targets. Nature 457:1133–1136.

Chen CC, Lu J, Yang R, Ding JB, Zuo Y (2018) Selective activation of parvalbumin interneurons prevents stress-induced synapse loss and perceptual defects. Mol Psychiatry 23:1614–1625.

Clemens AM, Lenschow C, Beed P, Li L, Sammons R, Naumann RK, Wang H, Schmitz D, Brecht M (2019) Estrus-Cycle Regulation of Cortical Inhibition. Curr Biol 29:605–615 e606.

Cowan RL, Wilson CJ (1994) Spontaneous firing patterns and axonal projections of single corticostriatal neurons in the rat medial agranular cortex. J Neurophysiol 71:17–32.

Delevich K, Piekarski D, Wilbrecht L (2019a) Neuroscience: Sex Hormones at Work in the Neocortex. Curr Biol 29:R122–R125.

Delevich K, Thomas AW, Wilbrecht L (2019b) Adolescence and “Late Blooming” Synapses of the Prefrontal Cortex. Cold Spring Harb Symp Quant Biol.

Delevich K, Hall C, Boivin JR, Piekarski D, Zhang Y, Wilbrecht L (2019c) Prepubertal gonadectomy reveals sex differences in approach-avoidance behavior in adult mice. bioRxiv.

Dembrow NC, Chitwood RA, Johnston D (2010) Projection-specific neuromodulation of medial prefrontal cortex neurons. J Neurosci 30:16922–16937.

DePasque S, Galvan A (2017) Frontostriatal development and probabilistic reinforcement learning during adolescence. Neurobiol Learn Mem 143:1–7.

DePoy LM, Shapiro LP, Kietzman HW, Roman KM, Gourley SL (2019) beta1-Integrins in the Developing Orbitofrontal Cortex Are Necessary for Expectancy Updating in Mice. J Neurosci 39:6644–6655.

Drzewiecki CM, Willing J, Juraska JM (2016) Synaptic number changes in the medial prefrontal cortex across adolescence in male and female rats: A role for pubertal onset. Synapse 70:361–368.

Furth D, Vaissiere T, Tzortzi O, Xuan Y, Martin A, Lazaridis I, Spigolon G, Fisone G, Tomer R, Deisseroth K, Carlen M, Miller CA, Rumbaugh G, Meletis K (2018) An interactive framework for whole-brain maps at cellular resolution. Nat Neurosci 21:139–149.

Gabbott PL, Warner TA, Jays PR, Salway P, Busby SJ (2005) Prefrontal cortex in the rat: projections to subcortical autonomic, motor, and limbic centers. J Comp Neurol 492:145–177.

Gentry RT, Wade GN (1976) Androgenic control of food intake and body weight in male rats. J Comp Physiol Psychol 90:18–25.

Gerfen CR, Paletzki R, Heintz N (2013) GENSAT BAC cre-recombinase driver lines to study the functional organization of cerebral cortical and basal ganglia circuits. Neuron 80:1368–1383.

Giedd JN, Blumenthal J, Jeffries NO, Castellanos FX, Liu H, Zijdenbos A, Paus T, Evans AC, Rapoport JL (1999) Brain development during childhood and adolescence: a longitudinal MRI study. Nat Neurosci 2:861–863.

Glantz LA, Gilmore JH, Hamer RM, Lieberman JA, Jarskog LF (2007) Synaptophysin and postsynaptic density protein 95 in the human prefrontal cortex from mid-gestation into early adulthood. Neuroscience 149:582–591.

Goddings AL, Beltz A, Peper JS, Crone EA, Braams BR (2019) Understanding the Role of Puberty in Structural and Functional Development of the Adolescent Brain. J Res Adolesc 29:32–53.

Gonzalez-Burgos G, Miyamae T, Pafundo DE, Yoshino H, Rotaru DC, Hoftman G, Datta D, Zhang Y, Hammond M, Sampson AR, Fish KN, Ermentrout GB, Lewis DA (2015) Functional Maturation of GABA Synapses During Postnatal Development of the Monkey Dorsolateral Prefrontal Cortex. Cereb Cortex 25:4076–4093.

Gourley SL, Olevska A, Warren MS, Taylor JR, Koleske AJ (2012) Arg kinase regulates prefrontal dendritic spine refinement and cocaine-induced plasticity. J Neurosci 32:2314–2323.

Ha GE, Cheong E (2017) Spike Frequency Adaptation in Neurons of the Central Nervous System. Exp Neurobiol 26:179–185.

Hammerslag LR, Gulley JM (2014) Age and sex differences in reward behavior in adolescent and adult rats. Dev Psychobiol 56:611–621.

Harris KD, Shepherd GM (2015) The neocortical circuit: themes and variations. Nat Neurosci 18:170–181.

Hart G, Bradfield LA, Balleine BW (2018a) Prefrontal Corticostriatal Disconnection Blocks the Acquisition of Goal-Directed Action. J Neurosci 38:1311–1322.

Hart G, Bradfield LA, Fok SY, Chieng B, Balleine BW (2018b) The Bilateral Prefronto-striatal Pathway Is Necessary for Learning New Goal-Directed Actions. Curr Biol 28:2218–2229 e2217.

Hattox AM, Nelson SB (2007) Layer V neurons in mouse cortex projecting to different targets have distinct physiological properties. J Neurophysiol 98:3330–3340.

Hintiryan H, Foster NN, Bowman I, Bay M, Song MY, Gou L, Yamashita S, Bienkowski MS, Zingg B, Zhu M, Yang XW, Shih JC, Toga AW, Dong HW (2016) The mouse cortico-striatal projectome. Nat Neurosci 19:1100–1114.

Hinton EA, Li DC, Allen AG, Gourley SL (2019) Social isolation in adolescence disrupts cortical development and goal-dependent decision making in adulthood, despite social reintegration. eNeuro.

Hodge RD et al. (2019) Conserved cell types with divergent features in human versus mouse cortex. Nature.

Holtmaat A, Bonhoeffer T, Chow DK, Chuckowree J, De Paola V, Hofer SB, Hubener M, Keck T, Knott G, Lee WC, Mostany R, Mrsic-Flogel TD, Nedivi E, Portera-Cailliau C, Svoboda K, Trachtenberg JT, Wilbrecht L (2009) Longterm, high-resolution imaging in the mouse neocortex through a chronic cranial window. Nat Protoc 4:1128–1144.

Holtmaat AJ, Trachtenberg JT, Wilbrecht L, Shepherd GM, Zhang X, Knott GW, Svoboda K (2005) Transient and persistent dendritic spines in the neocortex in vivo. Neuron 45:279–291.

Hooks BM, Papale AE, Paletzki RF, Feroze MW, Eastwood BS, Couey JJ, Winnubst J, Chandrashekar J, Gerfen CR (2018) Topographic precision in sensory and motor corticostriatal projections varies across cell type and cortical area. Nat Commun 9:3549.

Huizinga M, Dolan CV, van der Molen MW (2006) Age-related change in executive function: Developmental trends and a latent variable analysis. Neuropsychologia 44:2017–2036.

Hunnicutt BJ, Jongbloets BC, Birdsong WT, Gertz KJ, Zhong H, Mao T (2016) A comprehensive excitatory input map of the striatum reveals novel functional organization. Elife 5.

Huttenlocher PR (1979) Synaptic density in human frontal cortex – developmental changes and effects of aging. Brain Res 163:195–205.

Inoue T, Zakikhani M, David S, Algire C, Blouin MJ, Pollak M (2010) Effects of castration on insulin levels and glucose tolerance in the mouse differ from those in man. Prostate 70:1628–1635.

Jardi F, Laurent MR, Kim N, Khalil R, De Bundel D, Van Eeckhaut A, Van Helleputte L, Deboel L, Dubois V, Schollaert D, Decallonne B, Carmeliet G, Van den Bosch L, D’Hooge R, Claessens F, Vanderschueren D (2018) Testosterone boosts physical activity in male mice via dopaminergic pathways. Sci Rep 8:957.

Johnson C, Wilbrecht L (2011) Juvenile mice show greater flexibility in multiple choice reversal learning than adults. Dev Cogn Neuros-Neth 1:540–551.

Johnson CM, Loucks FA, Peckler H, Thomas AW, Janak PH, Wilbrecht L (2016) Long-range orbitofrontal and amygdala axons show divergent patterns of maturation in the frontal cortex across adolescence. Dev Cogn Neurosci 18:113–120.

Kaufman MT, Churchland MM, Ryu SI, Shenoy KV (2014) Cortical activity in the null space: permitting preparation without movement. Nat Neurosci 17:440–448.

Kiritani T, Wickersham IR, Seung HS, Shepherd GM (2012) Hierarchical connectivity and connection-specific dynamics in the corticospinal-corticostriatal microcircuit in mouse motor cortex. J Neurosci 32:4992–5001.

Kita T, Kita H (2012) The Subthalamic Nucleus Is One of Multiple Innervation Sites for Long-Range Corticofugal Axons: A Single-Axon Tracing Study in the Rat. Journal of Neuroscience 32:5990–5999.

Koss WA, Belden CE, Hristov AD, Juraska JM (2014) Dendritic remodeling in the adolescent medial prefrontal cortex and the basolateral amygdala of male and female rats. Synapse 68:61–72.

Kritzer MF (2002) Regional, laminar, and cellular distribution of immunoreactivity for ER alpha and ER beta in the cerebral cortex of hormonally intact, adult male and female rats. Cereb Cortex 12:116–128.

Krotkiewski M, Kral JG, Karlsson J (1980) Effects of castration and testosterone substitution on body composition and muscle metabolism in rats. Acta Physiol Scand 109:233–237.

Lake BB et al. (2016) Neuronal subtypes and diversity revealed by single-nucleus RNA sequencing of the human brain. Science 352:1586–1590.

Laubach M, Amarante LM, Swanson K, White SR (2018) What, If Anything, Is Rodent Prefrontal Cortex? eNeuro 5.

Laube C, Suleiman AB, Johnson M, Dahl RE, van den Bos W (2017) Dissociable effects of age and testosterone on adolescent impatience. Psychoneuroendocrinology 80:162–169.

Levesque M, Charara A, Gagnon S, Parent A, Deschenes M (1996) Corticostriatal projections from layer V cells in rat are collaterals of long-range corticofugal axons. Brain Res 709:311–315.

Li N, Chen TW, Guo ZV, Gerfen CR, Svoboda K (2015) A motor cortex circuit for motor planning and movement. Nature 519:51–56.

Mallya AP, Wang HD, Lee HNR, Deutch AY (2019) Microglial Pruning of Synapses in the Prefrontal Cortex During Adolescence. Cereb Cortex 29:1634–1643.

Markham JA, Mullins SE, Koenig JI (2013) Periadolescent maturation of the prefrontal cortex is sex-specific and is disrupted by prenatal stress. J Comp Neurol 521:1828–1843.

Master SL, Eckstein MK, Gotlieb N, Dahl R, Wilbrecht L, Collins AGE (2019) Distentangling the systems contributing to changes in learning during adolescence. bioRxiv.

Mitra S, Ghosh N, Sinha P, Chakrabarti N, Bhattacharyya A (2015) Alteration in Nuclear Factor-KappaB Pathway and Functionality of Estrogen via Receptors Promote Neuroinflammation in Frontal Cortex after 1-Methyl-4-Phe-nyl-1,2,3,6-Tetrahydropyridine Treatment. Sci Rep 5:13949.

Morishima M, Kawaguchi Y (2006) Recurrent connection patterns of corticostriatal pyramidal cells in frontal cortex. J Neurosci 26:4394–4405.

Naka A, Adesnik H (2016) Inhibitory Circuits in Cortical Layer 5. Front Neural Circuits 10:35.

Naneix F, Marchand AR, Di Scala G, Pape JR, Coutureau E (2012) Parallel maturation of goal-directed behavior and dopaminergic systems during adolescence. J Neurosci 32:16223–16232.

Nelson LR, Bulun SE (2001) Estrogen production and action. J Am Acad Dermatol 45:S116–124.

Neufang S, Specht K, Hausmann M, Gunturkun O, Herpertz-Dahlmann B, Fink GR, Konrad K (2009) Sex differences and the impact of steroid hormones on the developing human brain. Cereb Cortex 19:464–473.

Ng LHL, Huang Y, Han L, Chang RC, Chan YS, Lai CSW (2018) Ketamine and selective activation of parvalbumin interneurons inhibit stress-induced dendritic spine elimination. Transl Psychiatry 8:272.

Nguyen TV, McCracken J, Ducharme S, Botteron KN, Mahabir M, Johnson W, Israel M, Evans AC, Karama S, Brain Development Cooperative G (2013) Testosterone-related cortical maturation across childhood and adolescence. Cereb Cortex 23:1424–1432.

Oswald MJ, Tantirigama ML, Sonntag I, Hughes SM, Empson RM (2013) Diversity of layer 5 projection neurons in the mouse motor cortex. Front Cell Neurosci 7:174.

Pan WX, Mao T, Dudman JT (2010) Inputs to the dorsal striatum of the mouse reflect the parallel circuit architecture of the forebrain. Front Neuroanat 4:147.

Paolicelli RC, Bolasco G, Pagani F, Maggi L, Scianni M, Panzanelli P, Giustetto M, Ferreira TA, Guiducci E, Dumas L, Ragozzino D, Gross CT (2011) Synaptic pruning by microglia is necessary for normal brain development. Science 333:1456–1458.

Paus T, Keshavan M, Giedd JN (2008) Why do many psychiatric disorders emerge during adolescence? Nat Rev Neurosci 9:947–957.

Paus T, Nawaz-Khan I, Leonard G, Perron M, Pike GB, Pitiot A, Richer L, Susman E, Veillette S, Pausova Z (2010) Sexual dimorphism in the adolescent brain: Role of testosterone and androgen receptor in global and local volumes of grey and white matter. Horm Behav 57:63–75.

Peper JS, Schnack HG, Brouwer RM, Van Baal GC, Pjetri E, Szekely E, van Leeuwen M, van den Berg SM, Collins DL, Evans AC, Boomsma DI, Kahn RS, Hulshoff Pol HE (2009) Heritability of regional and global brain structure at the onset of puberty: a magnetic resonance imaging study in 9-year-old twin pairs. Hum Brain Mapp 30:2184–2196.

Petanjek Z, Judas M, Simic G, Rasin MR, Uylings HB, Rakic P, Kostovic I (2011) Extraordinary neoteny of synaptic spines in the human prefrontal cortex. Proc Natl Acad Sci U S A 108:13281–13286.

Piekarski DJ, Boivin JR, Wilbrecht L (2017a) Ovarian Hormones Organize the Maturation of Inhibitory Neurotransmission in the Frontal Cortex at Puberty Onset in Female Mice. Curr Biol 27:1735–1745 e1733.

Piekarski DJ, Johnson CM, Boivin JR, Thomas AW, Lin WC, Delevich K EMG, Wilbrecht L (2017b) Does puberty mark a transition in sensitive periods for plasticity in the associative neocortex? Brain Res 1654:123–144.

Porrero C, Rubio-Garrido P, Avendano C, Clasca F (2010) Mapping of fluorescent protein-expressing neurons and axon pathways in adult and developing Thy1-eYFP-H transgenic mice. Brain Res 1345:59–72.

Rakic P, Bourgeois JP, Eckenhoff MF, Zecevic N, Gold-man-Rakic PS (1986) Concurrent overproduction of synapses in diverse regions of the primate cerebral cortex. Science 232:232–235.

Reiner A, Jiao Y, Del Mar N, Laverghetta AV, Lei WL (2003) Differential morphology of pyramidal tract-type and intratelencephalically projecting-type corticostriatal neurons and their intrastriatal terminals in rats. J Comp Neurol 457:420–440.

Selemon LD (2013) A role for synaptic plasticity in the adolescent development of executive function. Transl Psychiatry 3:e238.

Shay DA, Vieira-Potter VJ, Rosenfeld CS (2018) Sexually Dimorphic Effects of Aromatase on Neurobehavioral Responses. Front Mol Neurosci 11:374.

Shepherd GM (2013) Corticostriatal connectivity and its role in disease. Nat Rev Neurosci 14:278–291.

Sierra A, Gottfried-Blackmore A, Milner TA, Mcewen BS, Bulloch K (2008) Steroid hormone receptor expression and function in microglia. Glia 56:659–674.

Sohur US, Padmanabhan HK, Kotchetkov IS, Menezes JR, Macklis JD (2014) Anatomic and molecular development of corticostriatal projection neurons in mice. Cereb Cortex 24:293–303.

Sowell ER, Thompson PM, Holmes CJ, Jernigan TL, Toga AW (1999) In vivo evidence for post-adolescent brain maturation in frontal and striatal regions. Nat Neurosci 2:859–861.

Spear LP (2000) The adolescent brain and age-related behavioral manifestations. Neurosci Biobehav Rev 24:417–463.

Sturman DA, Mandell DR, Moghaddam B (2010) Adolescents exhibit behavioral differences from adults during instrumental learning and extinction. Behav Neurosci 124:16–25.

Tirelli E, Laviola G, Adriani W (2003) Ontogenesis of behavioral sensitization and conditioned place preference induced by psychostimulants in laboratory rodents. Neurosci Biobehav Rev 27:163–178.

Turner RS, DeLong MR (2000) Corticostriatal activity in primary motor cortex of the macaque. J Neurosci 20:7096–7108.

Vandenberg A, Piekarski DJ, Caporale N, Munoz-Cuevas FJ, Wilbrecht L (2015) Adolescent maturation of inhibitory inputs onto cingulate cortex neurons is cell-type specific and TrkB dependent. Front Neural Circuits 9:5.

Vijayakumar N, Op de Macks Z, Shirtcliff EA, Pfeifer JH (2018) Puberty and the human brain: Insights into adolescent development. Neurosci Biobehav Rev 92:417–436.

Wallace M, Luine V, Arellanos A, Frankfurt M (2006) Ovariectomized rats show decreased recognition memory and spine density in the hippocampus and prefrontal cortex. Brain Res 1126:176–182.

Wickham H (2016) Ggplot2 : elegrant graphics for data analysis. In: Use R!, Second edition. Edition, p 1 online resource. Switzerland: Springer,.

Wilson CJ (1987) Morphology and synaptic connections of crossed corticostriatal neurons in the rat. J Comp Neurol 263:567–580.

Winnubst J et al. (2019) Reconstruction of 1,000 Projection Neurons Reveals New Cell Types and Organization of Long-Range Connectivity in the Mouse Brain. Cell.

Zuo Y, Lin A, Chang P, Gan WB (2005) Development of longterm dendritic spine stability in diverse regions of cerebral cortex. Neuron 46:181–189.

