## Supplementary Material for "Sex and pubertal status influence dendritic spine density onto frontal corticostriatal projection neurons"

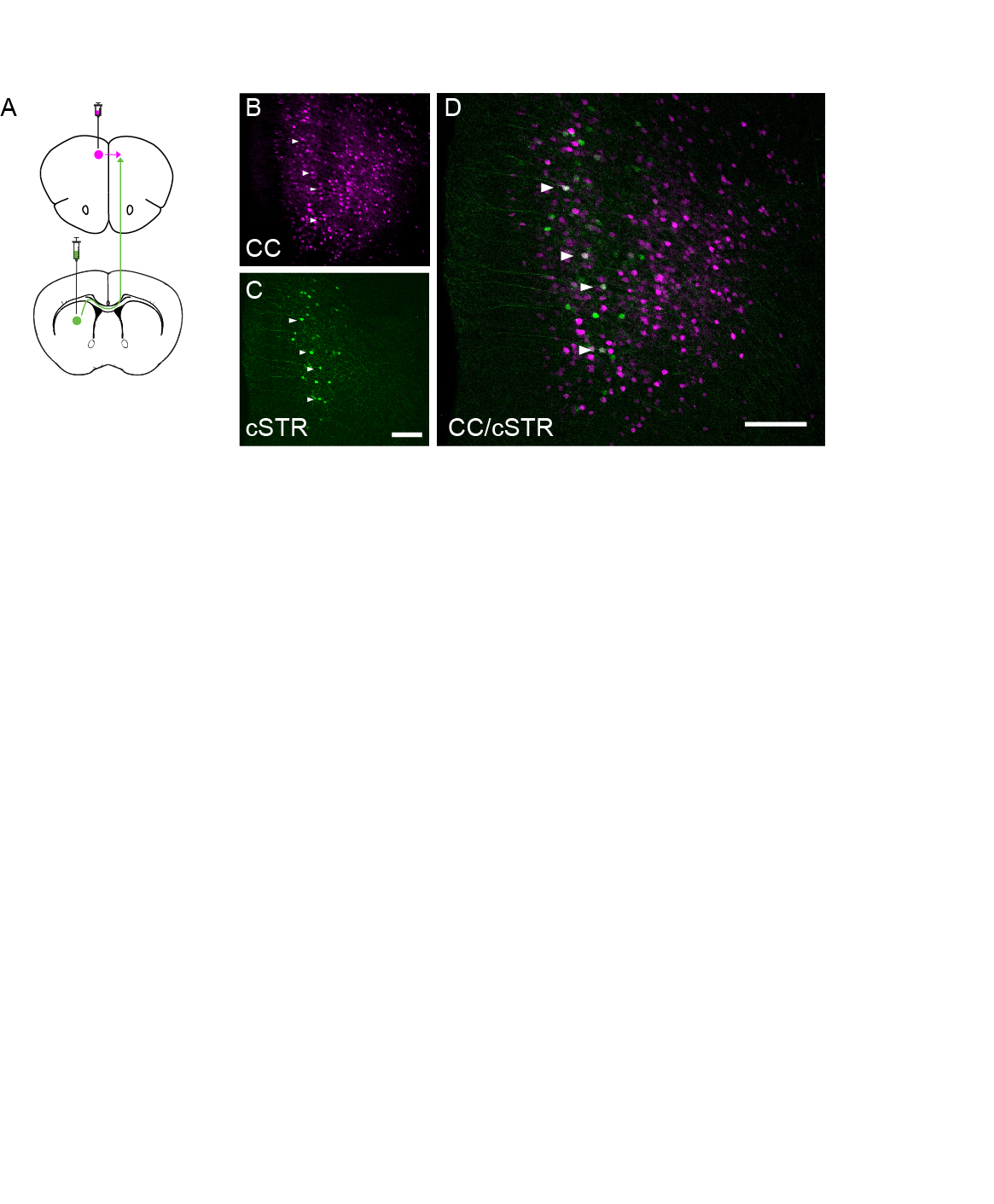

**Supplementary Figure 1: Subset of intratelencephalic (IT) cross-corticostriatal cells (cSTR) overlap with IT cortico-cortical (CC) cells in prelimbic cortex.** (A) Schematic of dual infusion of AAVretro-GFP into contralateral DMS and AAVretro-Tdtomato into contralateral prelimbic cortex. Images were acquired in the prelimbic cortex contralateral relative to the injection sites. (B) Labeled IT cortico-cortical cells in the prelimbic cortex. (C) Labeled IT corticostriatal cells in the prelimbic cortex (D) A subset of IT corticostriatal cells overlap with IT cortico-cortical cells in prelimbic cortex, as denoted by white arrows.

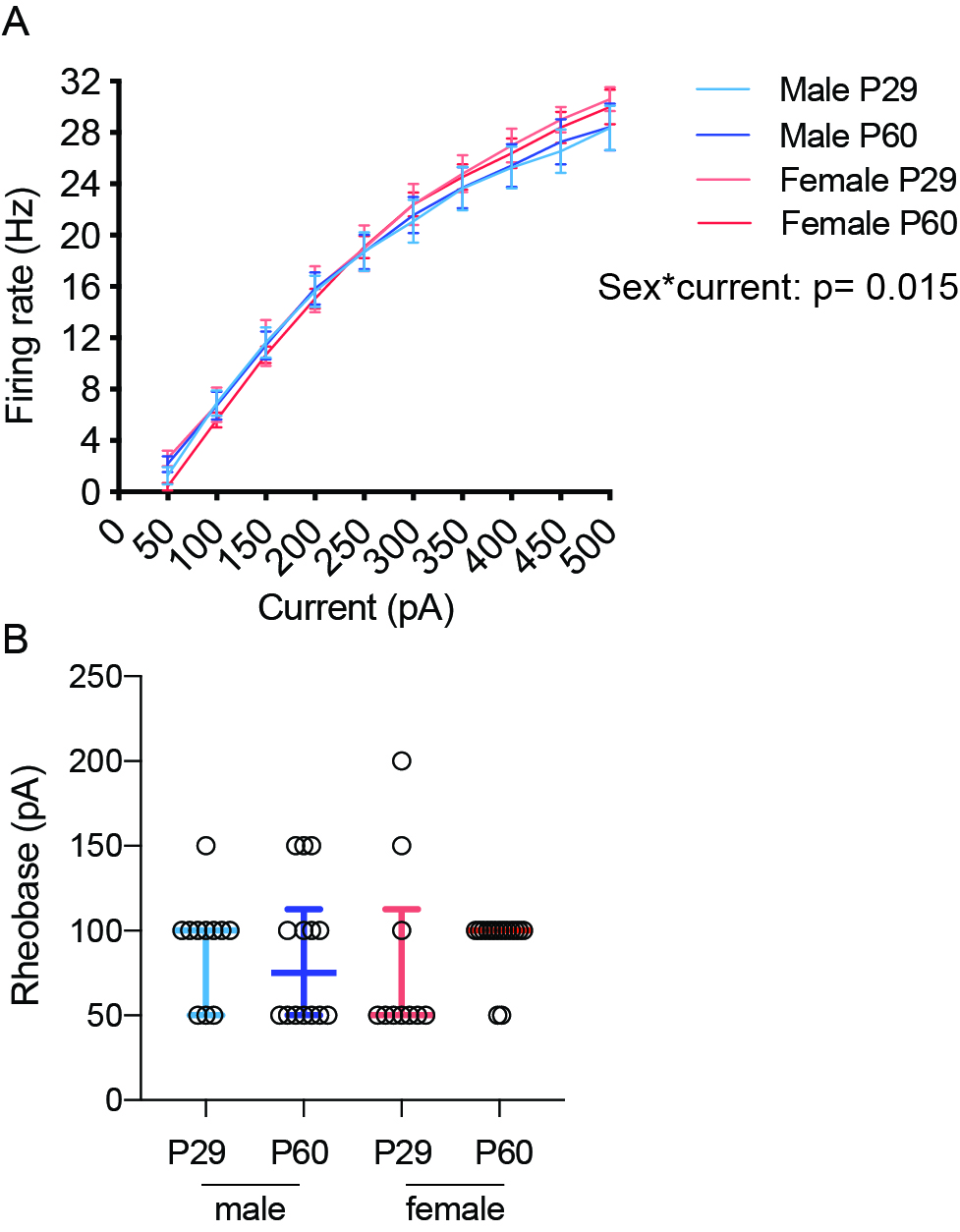

**Supplementary Figure 2: dmPFC cSTR neurons exhibit steeper input-output function compared to males.** (A) cSTR firing rate plotted as a function of square pulse current injection in juvenile and adult male and female mice. There was a significant interaction between sex and current injection [F(9, 414)= 2.317, p= 0.015 three-way repeated measures ANOVA] suggesting that females had a steeper relationship between current injection and firing rate. (B) There was no significant group effect on rheobase [H= 3.249, p= 0.35 Kruskal-Wallis one-way ANOVA].

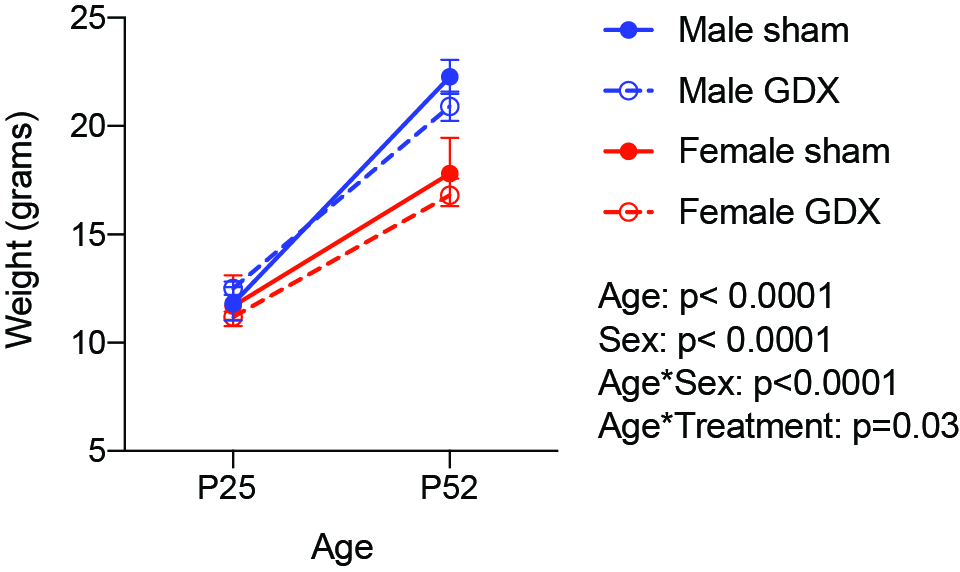

**Supplementary Figure 3: Sex and treatment effects on adolescent weight gain.** Weights of male and female mice were taken at postnatal day (P)25 when mice underwent sham or GDX surgery and at P52 when mice received viral transfusions into DMS for spine imaging experiments. There were significant main effects of age [F(1,18)= 687.6, p<0.0001] and sex [F(1,18)= 28.43, p<0.0001] on weight. In addition, there was a significant interaction between age and sex on weight [F(1,18)= 32.5, p<0.0001] indicating that males gained more weight over adolescence compared to females. Finally, there was a significant interaction between age and treatment (sham vs. GDX) [F(1,18)= 5.375, p= 0.0324], indicating that male and female mice who underwent GDX surgery gained less weight during adolescence relative to sham control mice. Data presented as mean ± SEM. Hypothesis testing performed using three-way repeated measures ANOVA.

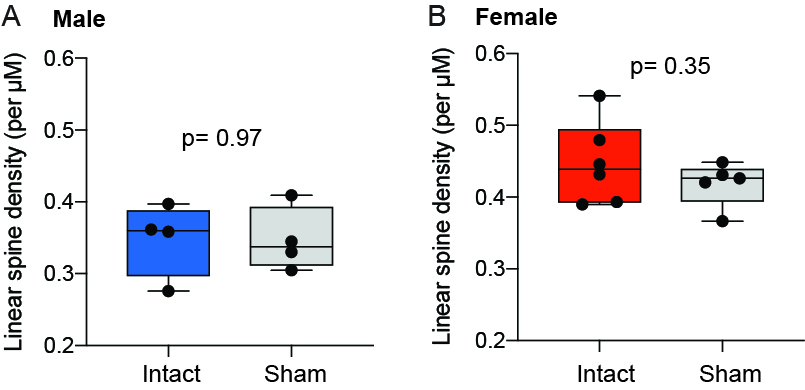

**Supplementary Figure 4: cSTR spine density is consistent between unmanipulated intact adult and sham surgery male and female mice.** (A) Male Intact (unmanipulated) vs. sham surgery at P25 did not differ in apical spine density at P60 (t_6_ = 0.037, p= 0.97). (B) Female Intact (unmanipulated) vs. sham surgery at P25 did not differ in apical spine density at P60 (t_9_ = 0.986, p= 0.35).

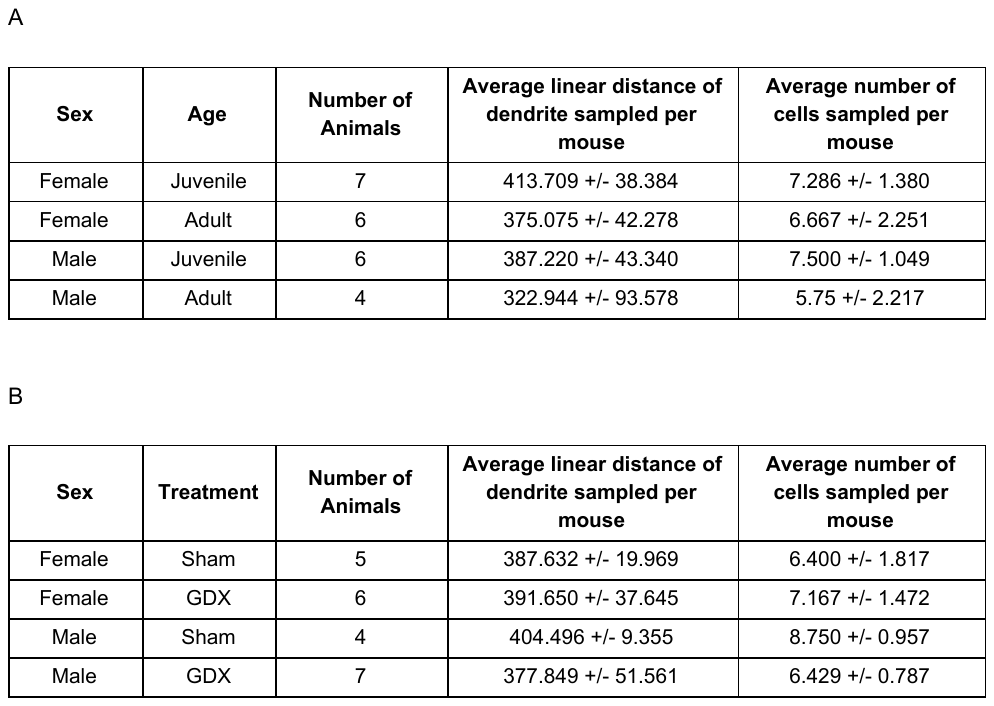

**Supplementary Table 1: Dendrite sampling across groups.**

Summary of group averages (mean ± SD) of dendrite distance (in µM) sampled per mouse for the developmental comparison (A) and pubertal status comparison (B). For each experimental group, the average linear distance of sampled dendrites (in µM) and the average number of cells sampled per mouse are presented.
